# Sustained coevolution of phage Lambda and *Escherichia coli* involves inner as well as outer membrane defenses and counter-defenses

**DOI:** 10.1101/2021.04.27.441663

**Authors:** Alita R. Burmeister, Rachel M. Sullivan, Jenna Gallie, Richard E. Lenski

**Affiliations:** Department of Microbiology and Molecular Genetics, Michigan State University, East Lansing, MI, USA; BEACON Center for the Study of Evolution in Action, Michigan State University, East Lansing, MI, USA; Department of Biology, University of Washington, Seattle, WA, USA; Department of Environmental Microbiology, Eawag, Dübendorf, Switzerland; Department of Environmental Systems Science, ETH Zürich, Zürich, Switzerland

## Abstract

Bacteria often evolve resistance to phage through the loss or modification of cell-surface receptors. In *Escherichia coli* and phage λ, such resistance can catalyze a coevolutionary arms race focused on host and phage structures that interact at the outer membrane. Here, we analyze another facet of this arms race involving interactions at the inner membrane, whereby *E. coli* evolves mutations in mannose permease-encoding genes *manY* and *manZ* that impair λ’s ability to eject its DNA into the cytoplasm. We show that these *man* mutants arose concurrently with the arms race at the outer membrane. We tested the hypothesis that λ evolved an additional counter-defense that allowed them to infect bacteria with deleted *man* genes. The deletions severely impaired the ancestral λ, but some evolved phage grew well on the deletion mutants, indicating they regained infectivity by evolving the ability to infect hosts independently of the mannose permease. This coevolutionary arms race fulfills the model of an inverse-gene-for-gene infection network. Taken together, the interactions at both the outer and inner membranes reveal that coevolutionary arms races can be richer and more complex than is often appreciated.

**IMPACT STATEMENT:** Laboratory studies of coevolution help us understand how host defenses and pathogen counter-defenses change over time, which is often essential for predicting the future dynamics of host-pathogen interactions. One particular model, termed “inverse-gene-for-gene” coevolution, predicts that coevolution proceeds through alternating steps, whereby hosts lose the features exploited by pathogens, and pathogens evolve to exploit alternative features. Using a classic model system in molecular biology, we describe the nature and timing of a previously overlooked step in the coevolution of *E. coli* and bacteriophage lambda. Our work demonstrates that this mode of coevolution can profoundly re-shape the interactions between bacteria and phage.

## INTRODUCTION

An issue of longstanding interest is whether the coevolution of bacteria and virulent (lytic) phages involves endless rounds of bacterial defenses and phage counter-defenses. Based on experiments in chemostats, Lenski and Levin (1) suggested that bacteria typically had the upper hand, as *Escherichia coli* often eventually evolved resistance by deleting or inactivating the phage’s specific receptor, which the phage could not readily overcome. This resistance did not imply the extinction of the phage, however, because it often reduced the bacteria’s competitiveness for resources. Instead, the typical outcome was coexistence of resistant and sensitive bacteria, with the latter more efficient at exploiting resources and thus able to sustain the phage’s persistence (2, 3). A study of cyanobacteria and their phages in the marine environment also supported this pattern (4). On the other hand, Lenski and Levin also pointed out that bacteria would lose the upper hand if the phage targeted a receptor that was essential for the bacteria to survive in their current environment. They cited then-recent work by Williams Smith & Huggins (5, 6), who showed they could successfully treat mice with otherwise lethal bacterial infections using a phage that specifically targeted a receptor required for the bacteria to colonize the mice. As the problem of bacterial resistance to antibiotics has grown, similar strategies are now being tested in which phage that specifically target drug-efflux pumps are deployed as therapeutic agents (7-9). In the meantime, yet other forms of bacteria-phage coevolution have been discovered, including CRISPR systems in bacteria and countermeasures to avoid these defenses in phage (10-13).

Another part of the argument that bacteria had the upper hand in the coevolutionary arms race depended on the idea that, while phages could often counter minor mutations in receptors, it was much more difficult for them to evolve the ability to use another receptor if the bacteria simply stopped producing the usual receptor (1). However, more recent work has shown that some host-phage pairs can undergo longer coevolutionary cycles involving defenses and counter-defenses at the outer membrane (14-16), and some phages can evolve to use new receptors even on a short time scale (17). This coevolutionary dynamic – in which hosts lose structures exploited by specific pathogens, and those pathogens evolve to exploit alternative structures – is called inverse-gene-for-gene (IGFG) coevolution (18-21). This IGFG framework is useful for representing changes in coevolving communities of bacteria and phage (Fig. 1). For example, if phage cannot evolve to exploit new features after bacteria have evolved resistance, then phage populations may be evolutionarily static (22, 23). Conversely, if phage exploit essential features of the bacteria that cannot be eliminated, then the host’s evolution is constrained and phage infectivity may remain elevated (6, 8). Our study builds on one such example of IGFG coevolution, in which it was discovered that populations of a virulent strain of phage λ often evolved the ability to use another outer-membrane receptor after coevolving *E. coli* reduced their expression of the receptor that the phage had initially exploited (17, 24).

**Fig. 1.**
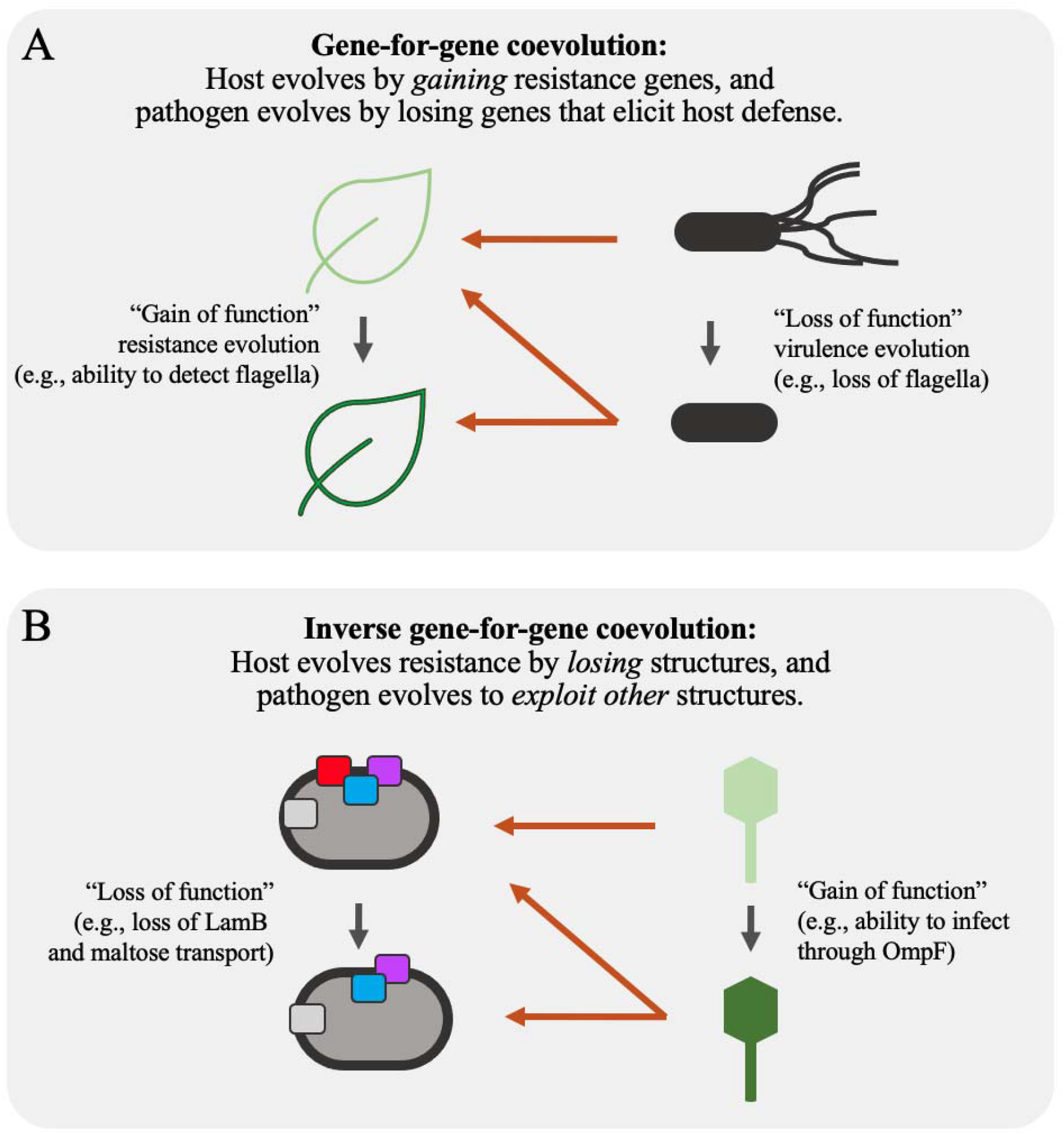
Genetic interaction networks during gene-for-gene (GFG) coevolution (panel A) and inverse-gene-for-gene (IGFG) coevolution (panel B). In both scenarios, host alleles affect selection on pathogen phenotypes, and pathogen alleles influence selection on host phenotypes. However, the two models have different implications for understanding historical coevolution and predicting future changes. During GFG coevolution, hosts evolve resistance by gaining resistance genes, and pathogens evolve by losing genes that elicit host defenses. GFG coevolution is common among plants and their bacterial pathogens; it may also occur in bacteria-phage interactions that involve restriction-modification and CRISPR defenses. During IGFG coevolution, pathogen infectivity requires the exploitation of specific host features, and resistance involves eliminating the exploited features. Unlike in the GFG model, host defenses in the IGFG model do not require pathogen recognition, and the pathogen’s evasion of host resistance does not require the loss of a defense elicitor.

Phage λ requires a two-step infection process to cross the outer and inner bacterial membranes (Fig. S1). The λ tail initiates infection at the outer membrane of the cell, where its J protein fibers adsorb to the bacterial protein LamB (25, 26). The tail proteins V and H allow λ to enter the periplasm and thereby interact with the mannose permease proteins (encoded by *manY* and *manZ*) in the inner membrane, which λ uses to eject its genome into the cytoplasm (27-30). Resistance to λ can occur by blocking λ’s entry at either the outer or inner membrane, with resistance mutations typically mapping to *lamB, lamB*’s positive regulator *malT* (25, 26), or the mannose permease genes (27, 28, 30) (Fig. S1). It has been shown that sensitive *E. coli* and lytic λ can coexist, along with resistant *E. coli* mutants, in both continuous (31) and batch culture regimes (17). Previous analysis of this coevolving system has revealed IGFG dynamics focused on outer membrane defenses and counter-defenses. That is, *E. coli* often first evolves *malT* mutations that reduce LamB expression, resulting in increased resistance to λ (17, 31, 32), and λ then regains infectivity through mutations in the *J* gene that increase its adsorption rate and fitness (31, 33). In some, but not all, experiments, specific sets of *J* mutations allow the novel exploitation of a second outer membrane protein, OmpF, catalyzing further evolution including mutations in the *ompF* gene (17, 34).

Despite extensive knowledge about the evolution of the initial (adsorption) and final (lysis) steps of λ infection of *E. coli*, much less is known about the evolution of the genetic networks during other stages of infection, including λ’s passage through the periplasmic space and the ejection of its DNA into the host cytoplasm. Meyer *et al*. (17) found that *E. coli* coevolving with λ often acquired mutations that impacted their ability to grow on mannose, which presumably were favored because they disrupt entry of the phage genome via the mannose permease. In this study, we examine how this coevolutionary arms race – previously focused on the cell’s outer membrane – also set off an arms race involving the host’s inner membrane, including the mechanism λ uses to eject its DNA through that membrane and into the bacteria’s cytoplasm.

## METHODS

### Bacteria and phage strains

Meyer *et al*. (17) founded 96 replicate cultures with *E. coli* B strain REL606 and lytic phage λ cI26, serially passaged the communities for 20 days, and froze mixed-community samples daily. Some of the phage populations evolved the ability to use the outer membrane protein OmpF as a receptor, some of the bacterial populations evolved mutations that affected mannose metabolism, and some communities changed in both respects. We obtained phage isolates from two of the populations (Table 1, Pop-A and Pop-B) that changed in both of these key respects; in each case, however, the isolates were taken four days before the phage had evolved the new ability to use the OmpF receptor (Table 1, Supplementary Material). *E. coli* K12 strains BW25113, JW1807, and JW1808 are from the Keio collection (35). REL606 Δ*manZ* was constructed using a two-step allelic exchange (Supplementary Material, Table S1, and Table S2).

**Table 1.** *E. coli* and phage λ strains used in this study.

| Strain | Description | Relevant Characteristics |
| --- | --- | --- |
| **Bacteria Clones:** |  |  |
| REL606 | *E. coli* B ancestor of coevolution experiment | *malT*<sup>+</sup>, *ompF*<sup>+</sup>, *manY*<sup>+</sup>, *manZ*<sup>+</sup> |
| REL606 $\Delta manZ$ | *manZ* deletion, derived from REL606<sup>a</sup> | $\Delta manZ$ |
| BW25113 | *E. coli* K12 parental strain of Keio collection | *malT*<sup>+</sup>, *ompF*<sup>+</sup>, *manY*<sup>+</sup>, *manZ*<sup>+</sup> |
| JW1807 | *manY* deletion in Keio collection | $\Delta manY$ |
| JW1808 | *manZ* deletion in Keio collection | $\Delta manZ$ |
| DH5α | Strain used for $\lambda$ plaque-based enumeration | *malT*<sup>+</sup>, *ompF*<sup>+</sup>, *manY*<sup>+</sup>, *manZ*<sup>+</sup> |
| **Phage Clones:** |  |  |
| cI26 | Lytic $\lambda$ ancestor of both phage populations | Requires *E. coli* LamB |
| λ-A | Evolved $\lambda$ isolate from Pop-A<sup>b</sup> on Day 8 (4 days before the population evolved to use OmpF) | Requires *E. coli* LamB |
| λ-B | Evolved $\lambda$ isolate from Pop-B<sup>b</sup> on Day 11 (4 days before the population evolved to use OmpF) | Requires *E. coli* LamB |

<sup>a</sup>This strain also has three mutations that have no known relevance to interactions with phage $\lambda$ (Supplementary Material). For construction methods, see Supplementary Material, Table S1, and Table S2.
<sup>b</sup>For simplicity, we have designated the source populations Pop-A and Pop-B. These correspond to population numbers D9 and G9 in the original experiment described by Meyer *et al.* (17).

### Phage growth assays

We measured the population growth of the ancestral and evolved phages under the same culture conditions as those in which the communities evolved (17) (Supplementary Material). The initial densities were ∼9 × 10^6^ cells per ml and ∼1 × 10^4^ phage per ml. We calculate the phage’s net population growth as the ratio of its final density after one day to its initial density; we show the resulting net growth on a log_10_-transformed scale. We enumerated the initial and final phage populations using dilution plating and soft-agar overlays (Supplementary Material). We performed 5 or 6 replicate assays for each phage-host combination shown in Figure 2.

**Fig. 2.**
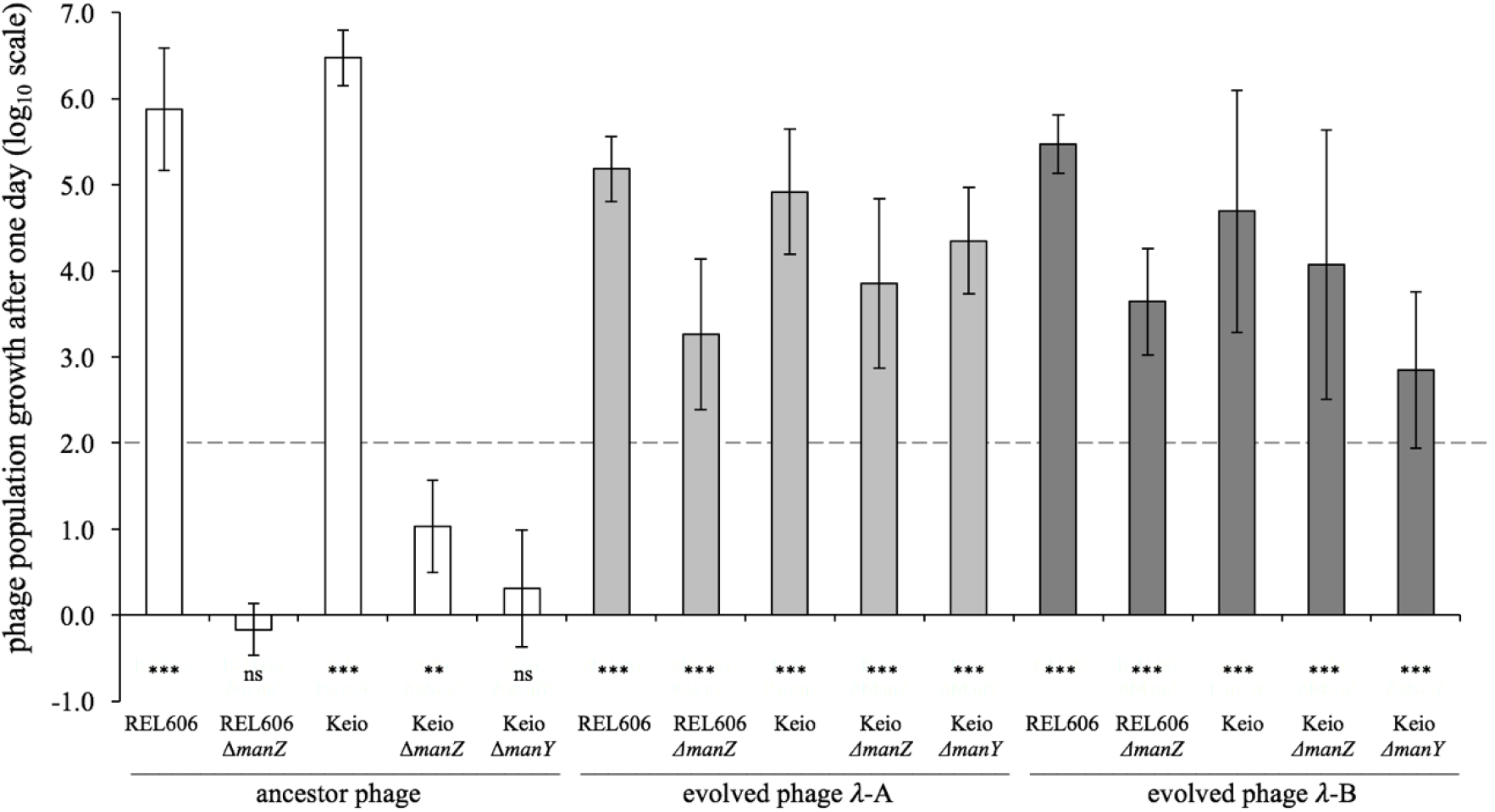
Net population growth of phage λ on wild type, Δ*manY*, and Δ*manZ* bacteria. Whether the phage could grow was assessed by performing one-tailed *t*-tests on the log_10_-transformed ratio of phage population densities at the start and end of a one-day cycle, with the null hypothesis of zero growth (***, *p* < 0.001; **, 0.001 < *p* < 0.01; ns, not significant, *p* > 0.05). Each test was based on 5 or 6 replicate assays. Phage isolates λ-A and λ-B evolved in a batch-culture regime with 100-fold dilution each day, and so 100-fold growth was required for their persistence; this break-even level is indicated by the dashed line.

### Frequency of mutants with altered mannose phenotypes

We estimated the frequency of bacteria with mutations affecting the mannose permease by plating from the time-series of frozen samples taken from populations Pop-A and Pop-B on tetrazolium mannose agar, as done previously (17). Mutants with reduced ability to metabolize mannose form deeply pigmented colonies that can be readily distinguished from those of the ancestral strain REL606, which forms light pink colonies on that medium.

#### Data accessibility

Data are available as Supplementary Datasets S1 (net population growth of phage λ on wild type and knockout bacteria) and S2 (temporal dynamics of *man* mutants in *E. coli* populations).

## RESULTS AND DISCUSSION

Our experiments focus on two independently coevolved communities of mixed *E. coli* and λ populations, designated Pop-A and Pop-B (17). Both λ populations evolved from a common ancestral phage (strain cI26). From each evolved population, we isolated a single phage clone: λ-A from Pop-A and λ-B from Pop-B (Table 1). Each clone was isolated 4 days before its population evolved the ability to use the OmpF receptor; hence, the phage clones were isolated on different days of the coevolution experiment performed by Meyer et al. (17).

To examine whether and how coevolution affected λ’s dependence on the ManY and ManZ proteins, we measured the population growth of the ancestral (cI26) and the two coevolved phage isolates (λ-A and λ-B) on bacterial strains with and without the *manY* and *manZ* genes (Table 1). Both the ancestral and evolved phage isolates grew well on bacterial strains with intact *manY* and *manZ* genes, including both the ancestral *E. coli* B strain, REL606, used in the coevolution experiment, and the K12 genetic background in which the Keio collection was made (Fig. 2, Table 1). Deletion of either the *manY* or *manZ* gene in either background severely reduced the ancestral phage’s population growth. In two cases (REL606 Δ*manZ* and Keio Δ*manY*), we saw no growth whatsoever in the ancestral phage (cI26) population after 24 hours; in the other case (Keio Δ*manZ*), the ancestral phage population increased ∼10-fold, but that was five orders of magnitude less than the increase on the same background with both mannose permease genes present. In striking contrast, both evolved phage isolates showed substantial growth on all three bacterial strains that lacked either the *manY* or *manZ* gene (Fig. 2). These results thus indicate an inverse-gene-for-gene coevolutionary interaction at the inner membrane. That is, the bacteria modified or lost the mannose permease, which the ancestral phage used to eject its genome into the cytoplasm, and the phage countered by evolving independence of that function.

To determine when the mutant mannose permease mutants arose in the two *E. coli* populations studied here, we plated frozen samples from the coevolution experiments on tetrazolium mannose agar, on which *man* mutants form pigmented colonies distinguishable from the wild type (Supplementary Material) (36). We are particularly interested in the timing of the appearance of the *man* mutants relative to two other steps in the coevolutionary arms race that were previously characterized: (i) the *malT* mutations that reduced the bacteria’s expression of LamB and thus the adsorption of the ancestral phage (33); and (ii) λ’s new ability to adsorb to OmpF as an alternative receptor (17). Our phage-growth data demonstrate that *manY* and *manZ* deletions confer substantial resistance even to the ancestral phage, which can use only the LamB surface receptor (Fig. 2). That result suggests the possibility that the *man* mutants could have arisen early in the coevolution experiments, perhaps alongside or even before the *malT* mutations that provided resistance at the outer membrane. However, time-course data show that the *man* alleles consistently reached high frequencies (above the detection limits, shown as gray dashed lines in Fig. 3) only after the fixation of the *malT* mutations, which occurred by day 8 in both populations studied here (17) (Fig. 3, Fig. 4, Table S3).

**Fig. 3.**
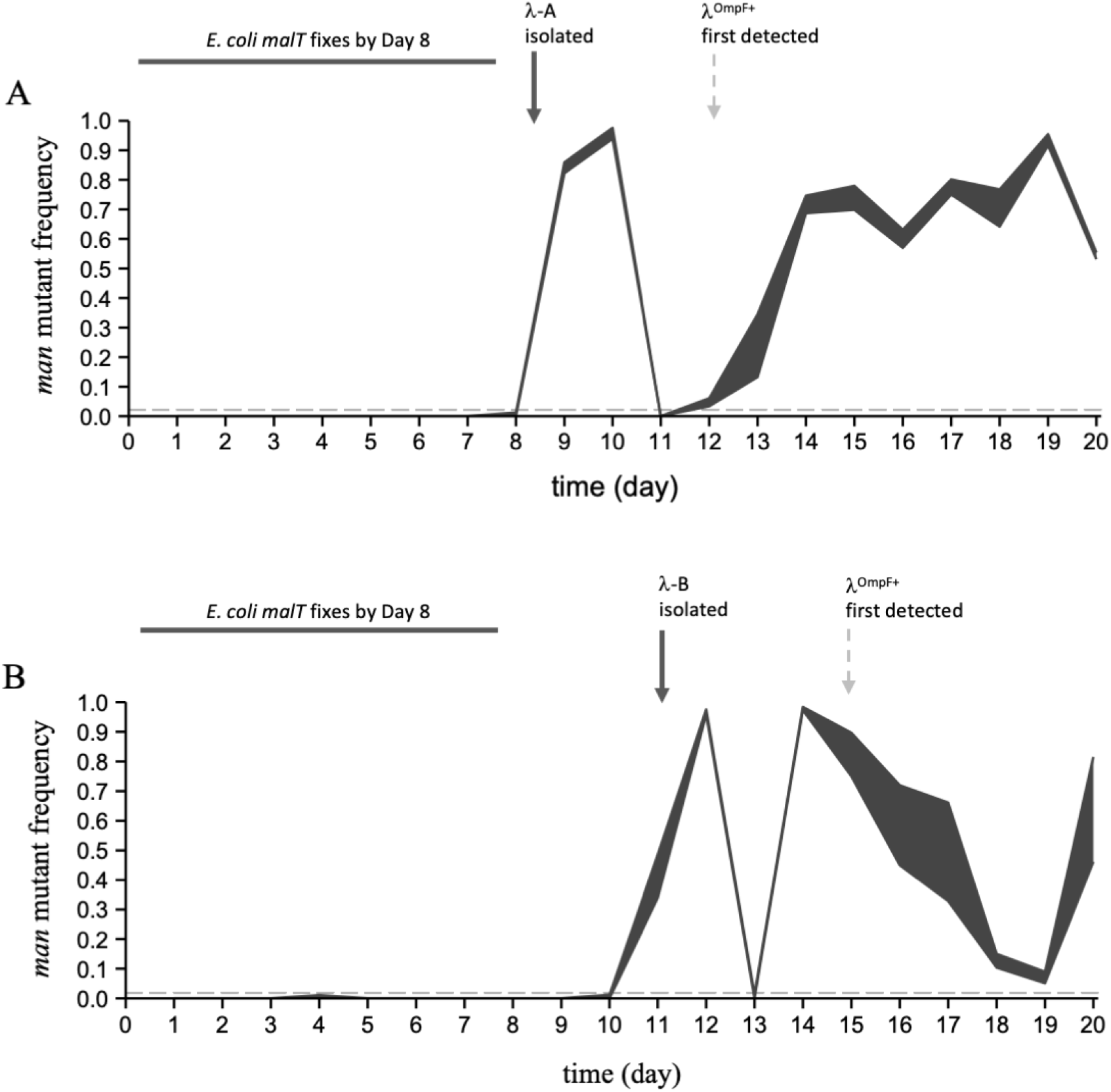
Temporal dynamics of *man* mutants in *E. coli* populations Pop-A (panel A) and Pop-B (panel B). Mutant *malT* alleles had already reached fixation in both populations by day 8 (17). Bacteria with *man* mutations, which confer resistance to the ancestral phage λ, rose to high frequencies and then declined sharply in abundance in both populations after day 8, but before λ had evolved to use the alternative receptor OmpF (timing indicated by vertical dashed arrows). These data imply that the *man* mutations evolved on *malT* mutant backgrounds, and that λ evolved independence of the mannose permease – causing the precipitous decline in the frequency of *man* mutants – before it evolved the ability to use OmpF. The shaded regions indicate the maximum and minimum frequencies of the *man* mutants based on analyzing two samples per population each day (mean *N* = 90 colonies tested per sample, minimum 29 colonies). The horizontal gray dashed lines show the approximate limit of detection of the *man* mutants (0.019 for panel A, 0.022 for panel B).

**Fig. 4.**
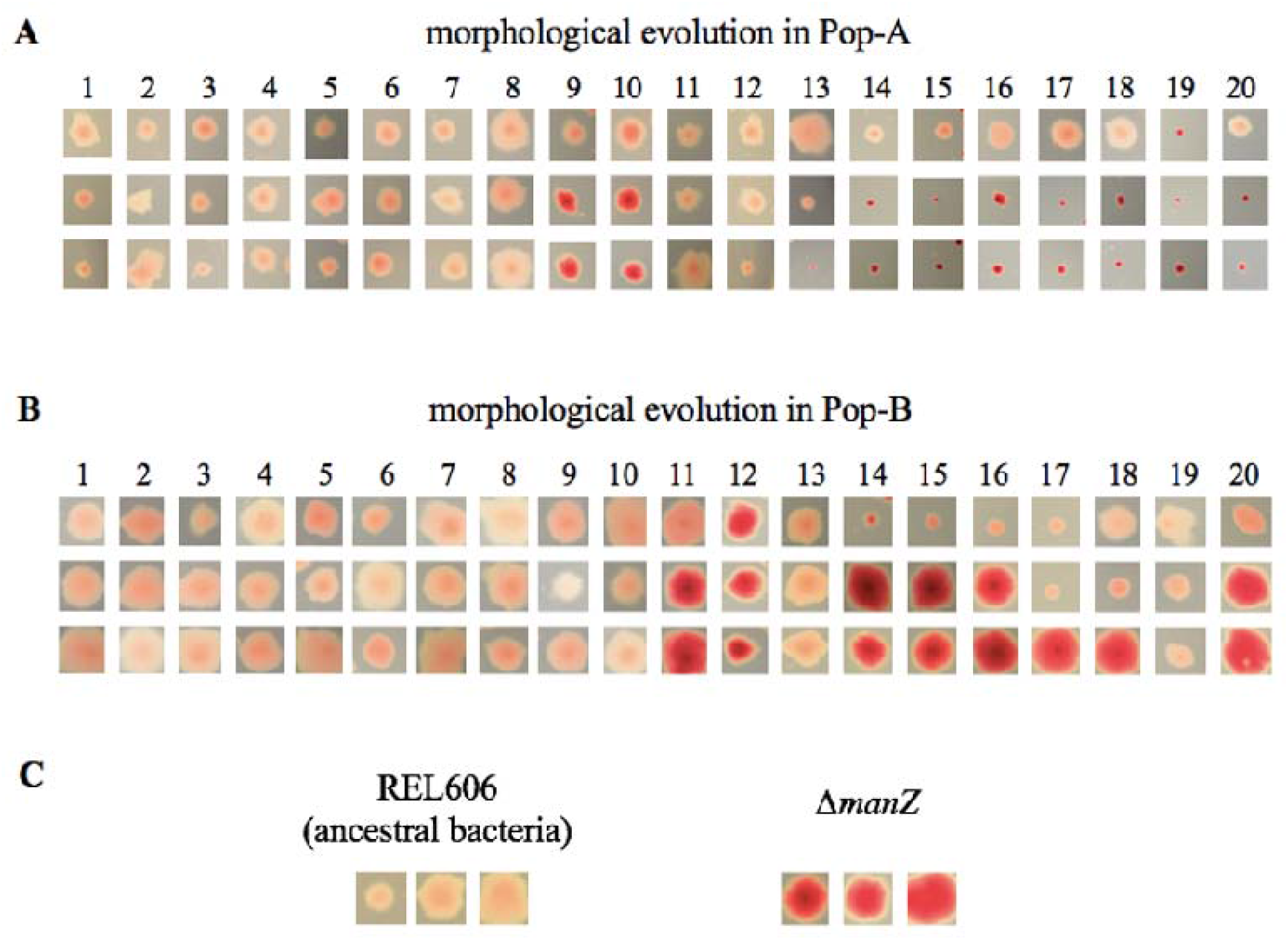
Evolution of *man*-related colony morphology on tetrazolium mannose agar. *E. coli* mutants with reduced ability to metabolize mannose form more deeply pigmented colonies than the wild type bacteria. Three representative colonies are shown for each sample from days 1-20 of two coevolution experiments. Representative colonies within a column are from the same agar plate and shown at the same magnification after incubation for 18-21 hours. Panel A: Pop-A. Panel B: Pop-B. Panel C: Comparison of wild type and Δ*manZ* bacteria in the same *E. coli* strain B genetic background.

These temporal data also show that the *man* mutations had nearly fixed in both bacterial populations (frequencies >95% on day 10 in Pop-A and on day 12 in Pop-B), but then the mutants sharply declined the next day. This reversal suggests these mutants were killed by phages that evolved independence of the mannose permease, and it is consistent with previous data showing that mutant *man* alleles rarely fixed in the bacterial populations (17). Meyer et al. (Fig. S2 in (17)) reported that the bacterial population densities remained high (∼2 × 10^9^ cells per ml, near the carrying capacity of the medium) throughout this period of the evolution experiment. Therefore, the mutant frequencies that we observed (Fig. 3) correspond to ∼4 × 10^7^ cells per ml (about 2% of the total population, the limit of detection in that assay) to almost 2 × 10^9^ cells per ml (the carrying capacity). With such large population sizes, any phage mutants that gained the ability to infect the *man* mutants would have access to a large number of hosts, and correspondingly, a large fitness benefit. The resulting growth of the *man*-independent phage population would drive the frequency of *man* mutants down, especially if the *man*-independent phages preferentially infected and killed the *man* mutants relative to other cells that retained the wild-type permease. Fitness costs associated with loss of the mannose permease may also have contributed to the reversal, although the costs of the resistance mutations are small compared to their benefit in the presence of phage (36).

In host Pop-A, variation in colony morphology further suggested that different *man* alleles were present before and after the sudden decline in the frequency of *man* mutants on day 11 (Fig. 4, Supplementary Material). The initial boom and bust of the mutant *man* alleles in both populations also occurred before the phage had evolved to use OmpF (Fig. 3, dashed arrows). Whether λ gained independence from the mannose permease by exploiting another inner membrane protein, and whether *E. coli* did (or could) respond by eliminating such a structure, are interesting questions for future work.

Our results are broadly consistent with genetic and molecular biology studies of λ host-range mutations. Scandella and Arber (30) isolated *E. coli* mutants that allowed phage adsorption to the cell envelope but interfered with ejection of the phage genome, thereby reducing infection success to a small fraction of that observed on wild-type cells. The responsible mutations were mapped to the mannose permease operon (27, 37), and λ mutants that could infect these mutant bacteria had mutations in phage genes *V* or *H* (38). Mutations in *V* and *H* have also been observed in another population in this study system (39). Williams *et al*. (37) found that, for *E. coli* strain K12, *manZ* is not strictly required for wild-type λ to eject its genome, and our results accord with that finding (Fig. 2, Keio background). However, our results suggest that λ cI26 does require *manZ* when infecting *E. coli* strain B, at least in the culture conditions that we used (Fig. 2, REL606 background). Alternatively, λ cI26 might occasionally infect and replicate in hosts without *manZ*, but at a rate that is offset by the decay or inactivation of free virus particles under these conditions (17, 33). In any case, the net population growth of the ancestral phage on either the Δ*manY* or Δ*manZ* bacteria is insufficient to offset the 100-fold daily dilutions (Fig. 2, dashed line) that took place during the coevolution experiment (17).

Taken together, our results imply that *E. coli* and λ coevolved in an inverse gene-for-gene manner (18) (Fig. 1). This coevolution involved two infection steps – crossing first the outer and then the inner membrane – and at least three, and probably four, distinct host features (Figs. 1, 5, and S1). *E. coli* evolved resistance to phage λ through the loss or alteration of maltose transport across the outer membrane (via mutations in *malT*) and mannose transport across the inner membrane (via mutations in *manY* or *manZ*), while λ evolved to exploit other *E. coli* features including another outer membrane protein (OmpF) and, presumably, some as yet unidentified, alternative inner membrane protein (shown as encoded by the hypothetical *imx* gene in Fig. 5). While our study addresses one particular bacteria-phage interaction in a simple laboratory setting, it illustrates the extent to which the resulting coevolutionary arms races can be richer and more complex than is often appreciated.

**Fig. 5.**
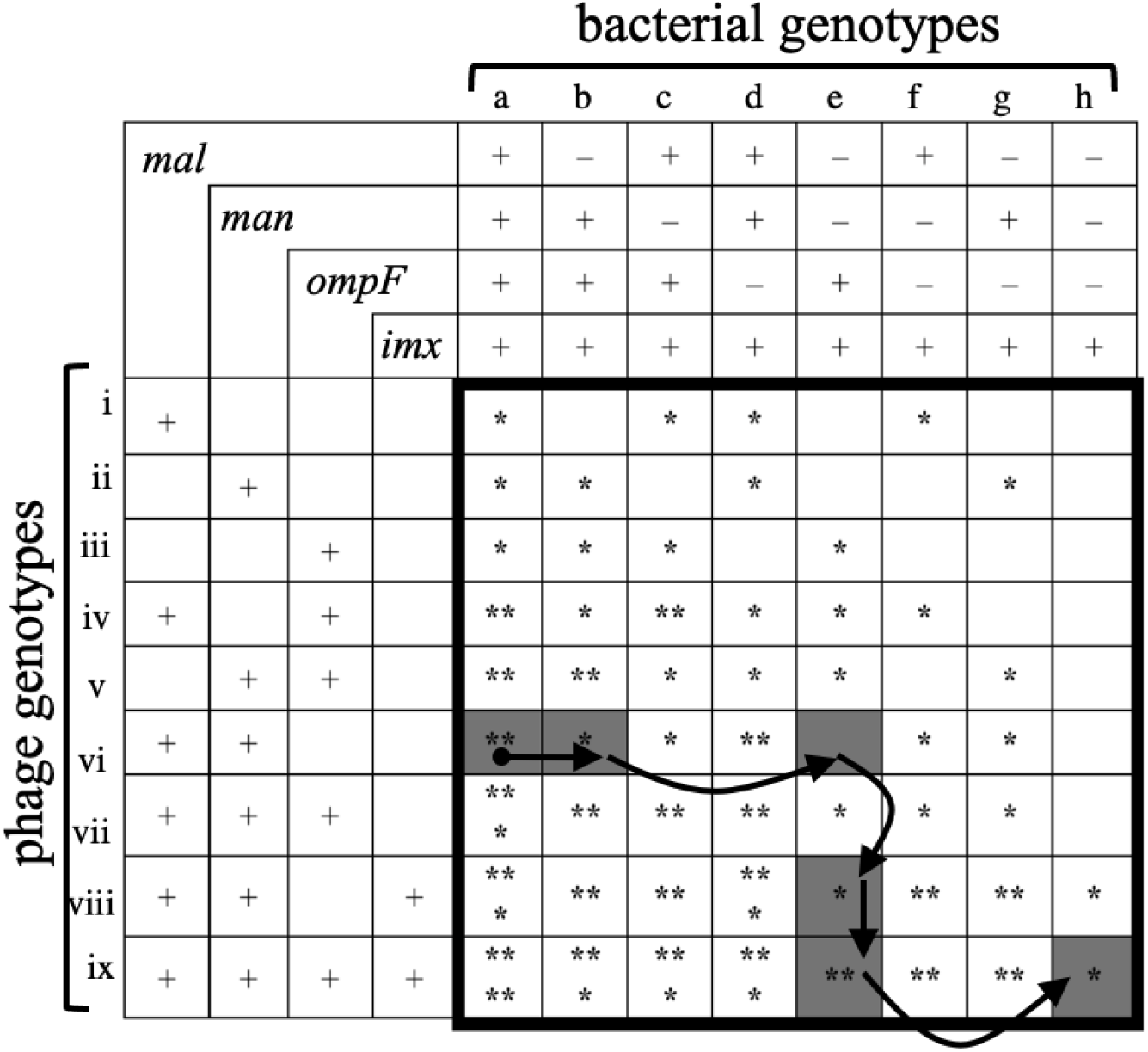
An inverse-gene-for-gene model showing the structure of the genetic network for coevolving *E. coli* and λ populations. Columns indicate bacterial genotypes with four exploitable features, and rows indicate λ genotypes that exploit those features: *mal*, maltose transport across the outer membrane; *man*, mannose transport across the inner membrane; *ompF*, glucose and electrolyte transport across the outer membrane; *imx*, a hypothetical inner <u>m</u>embrane feature that is exploited by λ that evolved independence of the mannose permease. The “+” symbol indicates that either the bacteria have the feature or the phage exploit the feature. The “–” symbol indicates the bacteria lack the feature, express it to a reduced degree, or otherwise modify it to minimize phage infection. Asterisks (*) indicate infectivity for each host-phage pair, with more asterisks indicating greater infectivity. Adaptive changes through the network can proceed by two types of moves: *E. coli* resistance (to the right across rows), and increased λ infectivity (downward across columns). The coevolving communities were founded by host genotype a and phage genotype vi (shown by the black circle). The communities analyzed in this study appear to have moved through the shaded nodes in five steps, as indicated by the arrows.

There are many alternative coevolutionary paths through an inverse-gene-for-gene network that has four features subject to host defenses and parasite counter-defenses (Fig. 5). This multiplicity of potential paths suggests that mutation and selection could drive replicate communities to different regions of the coevolutionary landscape, raising other interesting questions. How might different first-step resistance mutations affect the subsequent host-range evolution of the phage and the further evolution of host resistance? To what extent can IGFG systems continuously evolve host defenses and parasite counter-defenses? What is the effect of such prolonged coevolution for community diversity? Do communities become increasingly divergent as the coevolving populations follow different paths through the network, or might they eventually converge on the same phenotypic states after a period of divergence? How important are evolutionary innovations in opening new paths, relative to pleiotropic tradeoffs that may close off certain paths? Future work should investigate these and other questions about the coevolution of bacteria and phage and the structure of their genetic interaction networks.

## ACKNOWLEDGEMENTS

We thank Neerja Hajela for assistance in the lab; Justin Meyer for advice on experimental methods, sharing samples from the coevolution experiments, and valuable discussions; and Rohan Maddamsetti, Caroline Turner, and Mike Wiser for comments on the manuscript.

## AUTHOR CONTRIBUTIONS

A.R.B., R.M.S., and R.E.L. conceived the study. A.R.B., R.M.S., and J.G. performed the experiments. All authors analyzed the data and wrote the manuscript.

