## Supplemental Materials for "Sustained coevolution of phage Lambda and *Escherichia coli* involves inner as well as outer membrane defenses and counter-defenses"

**Supplemental Methods**

*Phage isolation from experimentally evolved communities:* We plated frozen samples taken from two experimentally evolved communities to obtain isolated plaques of phage clones λ-A and λ-B on lawns of *lamB*^+^ *E. coli* cells (Table 1). We confirmed that both of them require the outer membrane protein LamB for infection by observing that neither produces plaques on *lamB*^–^ *E. coli* lawns.

*Phage growth assays*: We measured phage growth in 10 ml of modified M9 (mM9) medium (M9 salts with 1 g/L magnesium sulfate, 1 g/L glucose, and 0.02% LB) in 50-ml glass flasks shaken at 120 rpm and kept at 37°C. To ensure independence of the replicate assays, the bacterial stocks for each replicate were each grown from a separate bacterial colony. We enumerated initial phage densities before adding cells and final phage densities without removing cells, which allowed us to quantify the final total density of viable phage including both infected hosts as well as free phage particles.

*Phage enumeration:* We estimated phage densities using dilution plating with soft-agar overlays. We mixed 100 µl of a diluted phage sample with 100 µl of a stationary-phase culture of DH5α cells grown in LB (~2 × 10^8^ cells) and 4 ml of reduced soft agar (rSA, 10 g/L tryptone, 1 g/L yeast extract, 8 g/L NaCl, 0.1% glucose, 2 mM of CaCl_2_, and 1.75 g/L agar). We then poured this mixture on top of a base plate of LB agar. This soft-agar formulation contains only 25% of the typical concentration of agar, which we found improves plaque visibility and estimated numbers for the experimentally evolved phage λ isolates.

*Deleting* manZ *from REL606*: We first constructed a deletion fragment, in which ~550 bp of DNA on either side of the *manZ* gene were seamlessly sewn together. This fragment was produced using splicing by overlap extension polymerase chain reaction (SOE-PCR) (1) with a high-fidelity DNA polymerase (Phusion, ThermoFisher). We then exchanged the deletion fragment with the homologous region of the REL606 chromosome in a spontaneous, RecA-dependent, two-step recombination process. We describe below the methods used for constructing the deletion fragment and for the allelic exchange.

Deletion fragment construction: Table S1 shows the primers used to construct the deletion fragment *via* SOE-PCR. The deletion fragment has 853 fewer bp than the corresponding wild type fragment; only the final 8 bp of the 861-bp *manZ* gene remain in the deletion mutant (Table S2). We confirmed the desired sequence of the constructed deletion fragment by ligation into pCR8/GW/TOPO (Invitrogen), transformation into One Shot TOP10 Chemically Competent *E. coli* Cells (Invitrogen), and Sanger sequencing (Macrogen USA). A sequence-confirmed deletion fragment was retrieved from pCR8/GW/TOPO by digestion with *Bgl*II, ligated into the *Bgl*II site of pKOV_unstuff (2, 3), and transformed into *E. coli* B REL606 made chemically competent (3). REL606 transformants carrying the pKOV_unstuff+deletion_fragment plasmid were selected on LB agar containing 20 μg/ml chloramphenicol (Cm) at 30˚C. At this temperature, replication from the temperature-sensitive M13 origin of replication in pKOV_unstuff is possible. At higher temperatures (43˚C and above), replication is prevented. We then grew an REL606 + vector transformant that was selected on LB + Cm at 30˚C in liquid LB + Cm (20 μg/ml), and we stored it in glycerol saline at –80˚C.

Allelic exchange, first step: The transformant described immediately above was selected on LB + Cm (20 μg/ml) agar at 43˚C in order to obtain a strain in which the vector was incorporated into the chromosome by RecA-mediated recombination (*i.e.*, first-step recombinants). The pKOV_unstuff vector (carrying *cat*, the Cm resistance gene) cannot replicate at 43˚C, and therefore only first-step recombinants can grow under these conditions. We isolated a first-step recombinant colony (Cm^R^ at 43˚C), grew the cells in liquid LB+Cm at 43˚C, and stored them in glycerol saline at –80˚C.

Allelic exchange, second step: The pKOV_unstuff vector can recombine out of the REL606 chromosome, taking with it either the *manZ* deletion fragment (leaving a genome indistinguishable from the original wild type) or the wild-type *manZ* fragment (leaving a *manZ* deletion mutant: Δ*manZ*), via the second RecA-mediated homologous recombination step. We used the *sacB* gene on the pKOV_unstuff vector to select for such rare spontaneous mutants. The *sacB* gene encodes a sucrose transporter, the expression of which causes death (as a result of increased osmotic pressure) when grown on LB medium deficient in NaCl and containing 5% sucrose (LB–NaCl + 5% sucrose).

The first-step recombinant described above was diluted (using LB–NaCl broth as dilutant) and plated onto LB–NaCl + 5% sucrose agar. We incubated plates at 30˚C (to promote expression of the plasmid-encoded *sacB*) for ~24 h. Several putative second-step recombinant colonies were selected, and we streaked them on fresh LB–NaCl + 5% sucrose agar plates to obtain clones. We grew a single colony from each purification streak in liquid LB (Miller’s formulation, containing 10 g/L NaCl) at 37˚C overnight (shaking), and stored the resulting cells in glycerol saline at –80˚C.

Confirming phenotypes and genotypes of the two-step recombinants: We used washed cells from each putative two-step recombinant strain as a PCR template. We used primers that amplify the REL606 *manZ* region, but are located outside of the constructed deletion fragment (manY_f/yobD_r; see Table S1). These primers give a product that is 2201 bp in REL606 and wild type double recombinants, but which is expected to be 1348 bp in Δ*manZ* recombinants (see Table S2). We checked the size of the PCR product for each putative two-step recombinant on an agarose gel. As expected, we observed a mixture of wild type-size and mutant-sized PCR products. We purified the remainder of the PCR product for a recombinant that gave rise to a mutant-sized PCR product, using the QIAquick PCR purification kit (QIAGEN), and we confirmed the sequence of the entire fragment by Sanger sequencing with manY_f and yboD_r (Macrogen USA) (Table S1). We then confirmed that this Δ*manZ* recombinant was sensitive to 20 μg/ml Cm in liquid medium (*i.e.*, it had lost the Cm resistance conferred by the pKOV_unstuff plasmid). We also confirmed that this Δ*manZ* recombinant was resistant to wild-type λ; the Δ*manZ* recombinant produced a confluent lawn when it was plated with the phage, whereas the parental REL606 strain showed no growth.

*Genome sequencing:* To sequence REL606 Δ*manZ*, we isolated genomic DNA using the Qiagen Genomic-tip 100/G kit (catalog number 10243). The sequencing library and sequence reads were generated by the Michigan State University Research Technology Support Facility on an Illumina MiSeq sequencer. We used the *breseq* pipeline (4) to align the sequence reads to an annotated reference genome of *E. coli* strain REL606.

**Supplemental Results and Discussion**

*Timing of* man *mutations:* Meyer *et al.* (5) reported that *malT* mutations had fixed in all of the bacterial populations by day 8 of their coevolution experiment, whereas mutations in *man* became common at later time points. We therefore expected that most or all of the *manYZ* mutations in our study would be present on *mal* mutant backgrounds. To confirm this expectation, we tested two clonal isolates from each population at each day on minimal maltose, minimal mannose, and permissive tetrazolium mannose agar plates. As expected, the *man* mutants were usually deficient in growth on the minimal maltose medium, indicating that they also carried *malT* mutations (Table S3). We previously reported these phenotypes, along with corresponding whole-genome sequences (6), showing that various mutations in *manY*, *manZ*, or both led to the altered tetrazolium mannose phenotype. We also know that few mutations reached appreciable frequencies in these *E. coli* populations (5-7). Most of the other mutations that reached substantial frequencies are in other genes known to confer phage resistance (*malT*, *ompF*, and *ompR*), none of which produce these changes on the tetrazolium mannose indicator agar (6). Therefore, we are confident from these multiple lines of evidence that these deeply pigmented colonies result from mutations in the *man* genes.

*Evolved mutations in* man *do not completely eliminate growth on mannose:* The isolates with mutations in *man* produced colonies that were, to varying degrees, red on tetrazolium mannose agar, which is indicative of diminished capacity to use mannose. Nonetheless, all of these isolates produced at least faint patches when spotted onto minimal mannose agar. This result indicates that the *man* mutations did not completely eliminate the ability to grow on mannose as a sole carbon source. In fact, the richly pigmented *man* isolates from bacterial Pop-B, as well as those from Pop-A before day 12, grew robustly on minimal mannose plates, whereas the Pop-A isolates from day 12 and later had severely reduced growth on minimal mannose plates.

The patch phenotypes on minimal mannose agar correlate with colony phenotypes on tetrazolium mannose plates as follows. The isolates that grew robustly on minimal mannose agar formed large magenta colonies on tetrazolium mannose plates; by contrast, those isolates with severely reduced growth on minimal mannose agar formed small red colonies on tetrazolium mannose plates. This result implies that some *man* mutations impair the mannose permease, and possibly also the entry of phage DNA, to a greater extent than do others.

*Mutations in the REL606* ∆manZ *knockout strain:* Sequencing the complete genome of the REL606 ∆*manZ* knockout strain revealed the expected *manZ* deletion (Δ853 bp starting at genome position 1,882,466) and three other mutations in genes not known to affect phage λ: (i) a 12,090-bp deletion beginning at genome position 787,879 that impacts an *E. coli* B-specific island of genes of unknown function including *ECB_00726* through *ECB_00739*; (ii) an IS-mediated insertion at genome position 3,656,171 in the intergenic region between *ECB_03412* and *yiaH*; and (iii) an IS-mediated deletion of 7,281 bp starting at position 3,894,997 that removes the ribose operon including genes *rbsD, rbsA, rbsC, rbsB, rbsK,* and *rbsR* along with part of the *yieO* gene*.*

**SUPPLEMENTAL FIGURES**

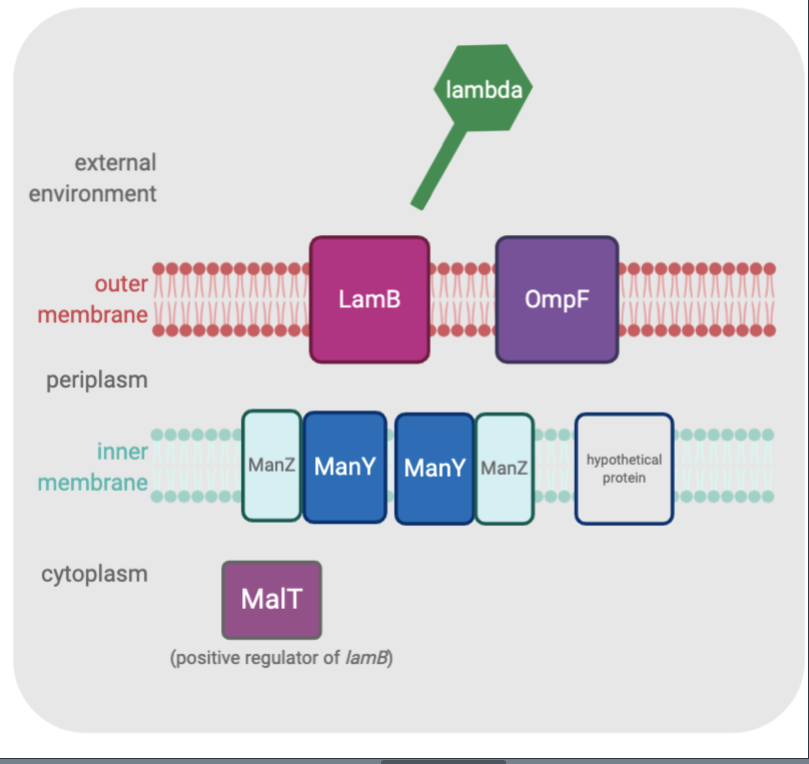

**Figure S1. Host features involved in λ infection.** Ancestral λ infects *E. coli* using the outer membrane protein LamB and the inner membrane mannose permease proteins ManY and ManZ. The host evolve resistance through losses of or modifications to LamB (or the *lamB* gene’s positive regulator MalT) and the ManYZ transmembrane channel. Meanwhile, λ evolves to become independent of those host features through use of OmpF and a hypothetical inner membrane feature (main text).

**SUPPLEMENTAL TABLES**

**Table S1.** Primers used in the construction and validation of the *manZ* deletion mutant in REL606. Capital letters denote the region annealing to the REL606 genome sequence, and small letters denote non-annealing primer sequence. Color highlighting corresponds to the primer nucleotide sequences shown in Table S2. Primers manY_f and yobD_r prime to sites in genes upstream and downstream of *manZ*, respectively. The first 4 primers listed were used to construct deletion fragments; the last 2 primers listed were used for final PCR and sequencing.

| **Primer Name** | **Sequence (5**'**🡪3**'**)^a^** |
| --- | --- |
| manZ_KO1 | gaagatctgttatcgcaggtcatcagagcattgg |
| manZ_KO2 | CTGTTAGTCCAGTTCGTTATCGAGATCG |
| manZ_KO3 | cgatctcgataacgaactggactaacagGACTGTAAGACTGTTGTACACTACCGG |
| manZ_KO4 | gaagatctGGAAGAGGTAATATAGCCTAAGCTATGTCTG |
| manY_f | gatgaacatcggtgctgcagttg |
| yobD_r | CCAAATGAGGGCGCAACCTTAAC |

^a^Underline denotes *Bgl*II restriction sites, which are used for the cloning process and are not included in the final, scar-free deletion.

**Table S2.** Wild-type and *manZ*-deletion sequences. Within each sequence, color highlighting indicates primer sites shown in Table S1. Text color indicates open reading frames of *manX* (blue), *manY* (red), *manZ* (green), and *yobD* (orange).

| **Sequence Name** | **Sequence (5**'**🡪3**'**)** |
| --- | --- |
| REL606 wild type chromosome, 1881377– 1884068 nt, forward strand | …gagcgtctcgttgaaggcggcgtgaaaatcacctctgttaacgtcggtggtatggcattccgtcagggtaaaacccaggtgaataacgcggtttcggttgatgaaaaagatatcgaggcgttcaagaaactgaatgcgcgcggtattgagctggaagtccgtaaggtttccaccgatccgaaactgaaaatgatggatctgatcagcaaaatcgataagtaacgtattgtgttgattatcactcagttttcacacttaagtcttacgtaaacaggagaagtacaatggagattaccactcttcaaattgtgctggtatttatcgtagcctgtatcgcaggtatgggatcaatcctcgatgaatttcagtttcaccgtccgctaatcgcgtgtaccctggtgggtatcgttcttggggatatgaaaaccggtattattatcggtggtacgctggaaatgatcgcgctgggctggatgaacatcggtgctgcagttgcgcctgacgccgctctggcttctatcatttctaccattctggttatcgcaggtcatcagagcattggtgcaggtatcgcactggcaatccctctggccgctgcgggccaggtactgaccatcatcgttcgtactattaccgttgctttccagcacgctgcggataaggctgctgataacggcaacctgacagcgatttcctggatccacgtttcttctctgttcctgcaagcaatgcgtgtggctattccggccgtcatcgttgcgctgtctgttggtaccagcgaagtacagaacatgctgaatgcgattccggaagtggtgaccaatggtctgaatatcgccggtggcatgatcgtggtggttggttatgcgatggttatcaacatgatgcgtgctggctacctgatgccgttcttctacctcggcttcgtaaccgcagcattcaccaactttaacctggttgctctgggtgtgattggtactgttatggcagtgctctacatccaacttagcccgaaatacaaccgcgtagccggtgcgcctgctcaggcagctggtaacaacgatctcgataacgaactggactaacaggtgagcgaaatggttgatacaactcaaactaccaccgagaaaaaactcactcaaagtgatattcgtggcgtcttcctgcgttctaacctcttccagggttcatggaacttcgaacgtatgcaggcactgggtttctgcttctctatggtaccggcaattcgtcgcctctaccctgagaacaacgaagctcgtaaacaagctattcgccgtcacctggagttctttaacacccagccgttcgtggctgcgccgattctcggcgtaaccctggcgctggaagaacagcgtgctaatggcgcagagatcgacgacggtgctatcaacggtatcaaagtcggtttgatggggccgctggctggtgtaggcgacccgatcttctggggaaccgtacgtccggtatttgcagcactgggtgccggtatcgcgatgagcggcagcctgttaggtccgctgctgttcttcatcctgtttaacctggtgcgtctggcaacccgttactacggcgtagcgtatggttactccaaaggtatcgatatcgttaaagatatgggtggtggcttcctgcaaaaactgacggaaggggcgtctatcctcggtctgtttgtcatgggggcattggttaacaagtggacacatgtcaacatcccgctggttgtctctcgcattactgaccagacgggcaaagaacacgttactactgtccagactattctggaccagttaatgccaggcctggtaccactgctgctgacctttgcttgtatgtggctactgcgcaaaaaagttaacccgctgtggatcatcgttggcttcttcgtcatcggtatcgctggttacgcttgcggcctgctgggactgtaagactgttgtacactaccggggccttttggccccgtttttttatctggaggattaatgacaatcacggacctggtactgattcttttcatcgccgcactcctggccttcgcgatctacgatcagttcatcatgccccgccgtaatggccccaccctgctggcaattcctttgctccggcgtggtcgcatcgatagcgttatcttcgtcggattgattgtcattcttatctataacaacgtcacgaatcatggtgcgttaataacgacatggttattaagcgcactggctctgatgggtttttatatattctggatccgcgttccgaagatcatctttaaacaaaaaggttttttcttcgccaatgtctggattgaatatagccgaatcaaagcgatgaacttgtcggaagatggcgtgttggtgatgcaattagaacagcgtcggctgttaatccgcgttcgaaatatcgacgatctggaaaaaatttataagcttctcgtttcaactcaataagttatgaatttagccaaagctatgtttagtgtatttttaataatcagacatagcttaggctatattacctcttcccttatttgttatttattttaacgtttcattgatatataaatctaaatgtaaatcgttatcaataaagcaatgaaataatatattccaacagttgttttatattctcaaaatatgttaaggttgcgccctcatttggggagtagccgatttccagattccggaaatgtacgtgt… |
| Δ*manZ* deletion fragment; primers manZ_KO2/3 sewn together, removing 853 bp | …gagcgtctcgttgaaggcggcgtgaaaatcacctctgttaacgtcggtggtatggcattccgtcagggtaaaacccaggtgaataacgcggtttcggttgatgaaaaagatatcgaggcgttcaagaaactgaatgcgcgcggtattgagctggaagtccgtaaggtttccaccgatccgaaactgaaaatgatggatctgatcagcaaaatcgataagtaacgtattgtgttgattatcactcagttttcacacttaagtcttacgtaaacaggagaagtacaatggagattaccactcttcaaattgtgctggtatttatcgtagcctgtatcgcaggtatgggatcaatcctcgatgaatttcagtttcaccgtccgctaatcgcgtgtaccctggtgggtatcgttcttggggatatgaaaaccggtattattatcggtggtacgctggaaatgatcgcgctgggctggatgaacatcggtgctgcagttgcgcctgacgccgctctggcttctatcatttctaccattctggttatcgcaggtcatcagagcattggtgcaggtatcgcactggcaatccctctggccgctgcgggccaggtactgaccatcatcgttcgtactattaccgttgctttccagcacgctgcggataaggctgctgataacggcaacctgacagcgatttcctggatccacgtttcttctctgttcctgcaagcaatgcgtgtggctattccggccgtcatcgttgcgctgtctgttggtaccagcgaagtacagaacatgctgaatgcgattccggaagtggtgaccaatggtctgaatatcgccggtggcatgatcgtggtggttggttatgcgatggttatcaacatgatgcgtgctggctacctgatgccgttcttctacctcggcttcgtaaccgcagcattcaccaactttaacctggttgctctgggtgtgattggtactgttatggcagtgctctacatccaacttagcccgaaatacaaccgcgtagccggtgcgcctgctcaggcagctggtaacaacgatctcgataacgaactggactaacaggactgtaagactgttgtacactaccggggccttttggccccgtttttttatctggaggattaatgacaatcacggacctggtactgattcttttcatcgccgcactcctggccttcgcgatctacgatcagttcatcatgccccgccgtaatggccccaccctgctggcaattcctttgctccggcgtggtcgcatcgatagcgttatcttcgtcggattgattgtcattcttatctataacaacgtcacgaatcatggtgcgttaataacgacatggttattaagcgcactggctctgatgggtttttatatattctggatccgcgttccgaagatcatctttaaacaaaaaggttttttcttcgccaatgtctggattgaatatagccgaatcaaagcgatgaacttgtcggaagatggcgtgttggtgatgcaattagaacagcgtcggctgttaatccgcgttcgaaatatcgacgatctggaaaaaatttataagcttctcgtttcaactcaataagttatgaatttagccaaagctatgtttagtgtatttttaataatcagacatagcttaggctatattacctcttcccttatttgttatttattttaacgtttcattgatatataaatctaaatgtaaatcgttatcaataaagcaatgaaataatatattccaacagttgttttatattctcaaaatatgttaaggttgcgccctcatttggggagtagccgatttccagattccggaaatgtacgtgt… |

**Table S3.** Almost all *man* mutants arose in genetic backgrounds that also evolved to be deficient in maltose metabolism. Colonies from samples taken on each day of the evolution experiment were streaked on tetrazolium mannose agar. For each day that yielded red colony phenotypes (indicating *man* mutations), two colonies were patched on minimal maltose, minimal mannose, and permissive tetrazolium arabinose agar plates. Most *man* mutants had growth deficiencies on minimal maltose agar. Some *man* mutants from Pop-A, and all of them from Pop-B, were nonetheless able to grow on minimal mannose agar. Therefore, we used the tetrazolium mannose phenotype for identifying the *man* mutant phenotype.

|  | **Colony 1 patches** | | | **Colony 2 patches** | | |
| --- | --- | --- | --- | --- | --- | --- |
| **Day** | **Minimal maltose** | **Minimal mannose** | **Tetrazolium arabinose** | **Minimal maltose** | **Minimal mannose** | **Tetrazolium arabinose** |
| *Pop-A* |  |  |  |  |  |  |
| 9 | – | + | + | + | + | + |
| 10 | – | + | + | – | + | + |
| 12 | – | – | + | – | – | + |
| 13 | – | – | + | – | + | + |
| 14 | – | – | + | – | + | + |
| 15 | – | – | + | – | – | + |
| 16 | – | – | + | – | – | + |
| 17 | – | – | + | – | – | + |
| 18 | – | – | + | – | – | + |
| 19 | – | – | + | – | – | + |
| 20 | – | – | + | – | – | + |
| *Pop-B* |  |  |  |  |  |  |
| 11 | + | + | + | – | + | + |
| 12 | – | + | + | – | + | + |
| 14 | – | + | + | – | + | + |
| 15 | – | + | + | – | + | + |
| 16 | – | + | + | – | + | + |
| 17 | – | + | + | – | + | + |
| 18 | – | + | + | – | + | + |
| 19 | – | + | + | – | + | + |
| 20 | – | + | + | – | + | + |
